# Controlling septum thickness by a large protein ring

**DOI:** 10.1101/771139

**Authors:** Michaela Wenzel, Ilkay N. Celik Gulsoy, Yongqiang Gao, Joost Willemse, Mariska G. M. van Rosmalen, Gijs J. L. Wuite, Wouter H. Roos, Leendert W. Hamoen

## Abstract

Gram-positive bacteria divide by forming a thick cross wall. How the thickness of this septal wall is controlled is unknown. In this type of bacteria, the key cell division protein FtsZ is anchored to the cell membrane by two proteins, FtsA and SepF. We have isolated SepF homologues from different bacterial species and found that they all polymerize into large protein rings with diameters varying from 19 to 41 nm. Importantly, these values correlated well with the thickness of their septa. To test whether ring diameter determines septal thickness, we tried to construct different SepF chimeras with the purpose to manipulate the diameter of the SepF protein ring. This was indeed possible and confirmed that the conserved core domain of SepF determines ring diameter. Importantly, when SepF chimeras with a smaller diameter were expressed in the bacterial host *Bacillus subtilis*, the thickness of its septa also became smaller. These results strongly support a model in which septal thickness is controlled by curved molecular clamps formed by SepF polymers attached to the leading edge of nascent septa. This also implies that the intrinsic shape of a protein polymer can function as a mould to shape the cell wall.

**Significance Statement:** Many bacteria form a thick cell wall and divide by forming a cross wall. How they control the thickness of their cell wall and cross wall is unknown. In this study we show that in these bacteria the cell division protein SepF forms very large protein rings with diameters that correspond to the diameter of their cross walls. Importantly, when we reduced the diameter of SepF rings in the bacterial host *Bacillus subtilis* the cross wall also became thinner. These results provide strong evidence that a large protein ring can function as a mould to control the thickness of the cell wall that divides these bacterial cells.

## Introduction

The hallmark of Gram-positive bacteria is their thick cell wall composed of multiple layers of peptidoglycan. They divide by synthesizing a crosswall in between the newly formed daughter cells, and the thickness of the nascent division septum approaches that of the lateral cell wall. How Gram-positive bacteria regulate the thickness of their division septum is not known.

Bacterial cell division is accomplished by a complex multi-protein machinery called the divisome. Assembly of the divisome begins with polymerization of the tubulin homologue FtsZ at midcell into a ring-like configuration, the so called Z-ring (1). This structure forms a scaffold for the late cell division proteins that are responsible for synthesis of the dividing septal wall (2). Several cell division proteins support the formation of the Z-ring, and a key step is the anchoring of FtsZ polymers to the cell membrane. This is achieved by the conserved peripheral membrane proteins FtsA and SepF. Both proteins directly interact with FtsZ and use an amphipathic α-helix to bind the cell membrane (3, 4). FtsA can be found in both Gram-positive and Gram-negative bacteria, whereas SepF is widely conserved in Gram-positive and Cyanobacteria, but has no known homologue in Gram-negatives (5, 6). Other Z-ring proteins are the conserved protein ZapA, which interlinks FtsZ polymers (7), and the bitopic transmembrane proteins EzrA (Gram-positives) and ZipA (Gram-negatives) (8, 9). Once the Z-ring is assembled, the late cell division proteins arrive. These conserved transmembrane proteins form a complex comprising the peptidoglycan glycosyltransferase FtsW (10), the transpeptidase Pbp2B (FtsI in Gram-negatives) (11, 12), and the heterotrimeric complex composed of FtsL, DivIC and DivIB (FtsL, FtsB and FtsQ in Gram-negatives, respectively) (13, 14). It is assumed that the latter three proteins regulate the assembly of late cell division proteins, although it is not yet clear how the late proteins are recruited to the Z-ring in Gram-positive bacteria.

Some bacteria, such as the Gram-positive model organism *Bacillus subtilis*, contain both FtsA and SepF, and this organism needs only one of them for Z-ring formation. However, the absence of SepF results in highly deformed septa, which is not the case when FtsA is absent (6). This indicates that SepF must have an additional function related to septum formation. A curious property of purified *B. subtilis* SepF is that it forms large ring structures with an inner diameter of 40 nm (15). Based on the SepF crystal structure, these rings must encompass at least 80 to 100 SepF molecules (15). *In vitro*, these protein rings are able to bundle FtsZ polymers into very long microtubule-like structures with SepF rings stacked perpendicularly to the FtsZ polymers (16). However, such microtubular structures have never been observed in bacteria, and later studies showed that the membrane-binding amphipathic α-helix of SepF is likely located inside the ring, which seems to rule out ring formation *in vivo* (15).

Interestingly, the inner diameter of SepF rings is about the same size as the thickness of the septal wall (43 nm). We wondered whether this relationship is relevant, and if so, whether SepF rings might actually control the thickness of the septum. To examine this, we purified SepF from different Gram-positive bacteria and found that all these proteins bind to lipid membranes and form large protein rings, albeit with different diameters. Importantly, also in these organisms there was a correlation between SepF ring diameter and septum thickness. To confirm that the SepF ring diameter determines septal width, we expressed SepF chimeras with smaller diameters in *B. subtilis*. Indeed, this reduced the thickness of septa. These results provide strong evidence that Gram-positive bacteria regulate the thickness of their septal wall by the strong curvature of SepF polymers at the leading edge of nascent septa. This also implies that the intrinsic form of a protein polymer can function as a mold that can shape a cell wall.

## Results

### SepF rings and tubules

Purified *B. subtilis* SepF forms large protein rings when observed with transmission electron microscopy (TEM) using negative staining with uranyl acetate (15). To confirm these findings with an independent method, we examined whether the protein can form rings under more physiological conditions using atomic force microscopy (AFM). It appeared that under these conditions purified *B. subtilis* SepF also forms rings with diameters of 41 nm (Fig. 1). Rings were 2 to 4 nm high (Fig. 1C, D), which corresponds to a single or double molecule stacking, since the crystal packing has shown that SepF polymers can be either 1.5 or 2.6 nm wide, depending on the orientation of SepF dimers (15).

**Fig. 1.**
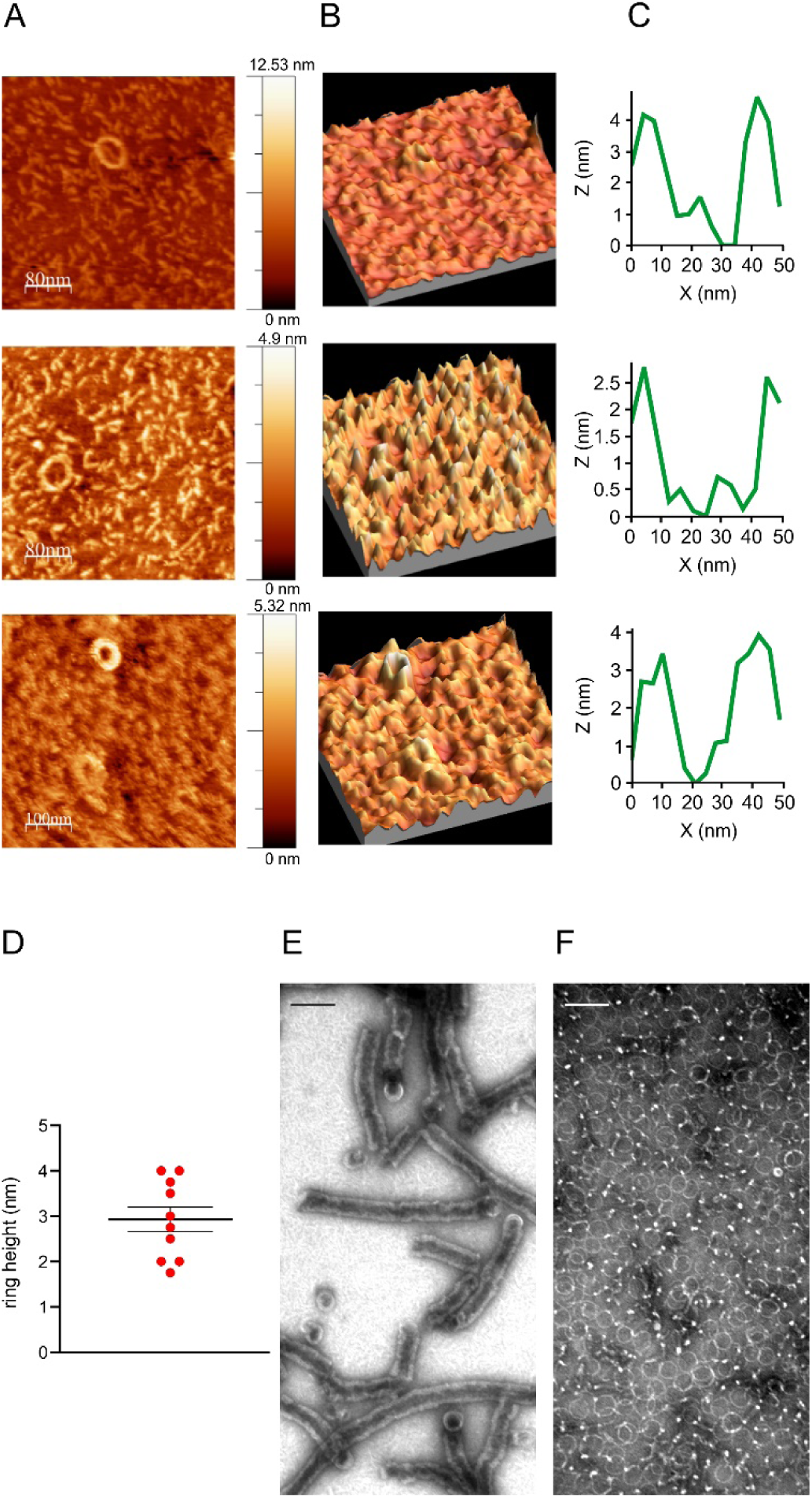
SepF forms rings and tubules. **(A)** Plane AFM images of three different SepF rings imaged in buffer solution. **(B)** 3D projections of the same images. **(C)** Height measurements of the individual SepF rings shown in A and B. **(D)** Height measurements of SepF rings derived from AFM data. **(E)** TEM image of SepF stack and tubules after cleavage from MBP. **(F)** TEM image of SepF rings after ion-exchange chromatography. Scale bars 100 nm.

Previously, SepF was purified as a maltose-binding protein (MBP) fusion followed by proteolytic cleavage and removal of MBP using ion-exchange chromatography (15, 16). We noticed that after cleavage, SepF precipitates, possibly due to the presence of calcium in the digestion buffer. After resuspension in calcium-free buffer, SepF formed stacks and tubules with diameters corresponding to that of SepF rings when observed by TEM (Fig. 1E). After ion-exchange chromatography only rings are found (Fig. 1F). These findings, together with the AFM data, show that the circular polymerization of *B. subtilis* SepF is a robust feature that may also occur *in vivo*.

### SepF from other species

To determine whether ring formation is a conserved feature of SepF, we purified the protein from different organisms. *Bacillus cereus* is an important food-spoiling bacterium (17, 18) and the causative agent of rainforest anthrax (19–21) and was chosen as a close relative of *B. subtilis*. We further selected *Clostridium perfringens*, *Mycobacterium tuberculosis*, and *Streptococcus pneumoniae,* all of which are important human pathogens. *S. pneumoniae* differs from the rest since it forms cocci instead of rods, and *M. tuberculosis* is one of the bacterial species that lack an FtsA homologue. An amino acid sequence alignment of the different SepF homologues is shown in Fig. 2A. The central domain of the protein, which contains the FtsZ binding site (15), is particularly conserved. All proteins were successfully purified as MBP fusions. *C. perfringens, M. tuberculosis,* and *S. pneumoniae* SepF precipitated after proteolytic cleavage from MBP and were collected by centrifugation, whereas *B. cereus* SepF remained largely soluble and was isolated by ion-exchange chromatography.

**Fig. 2.**
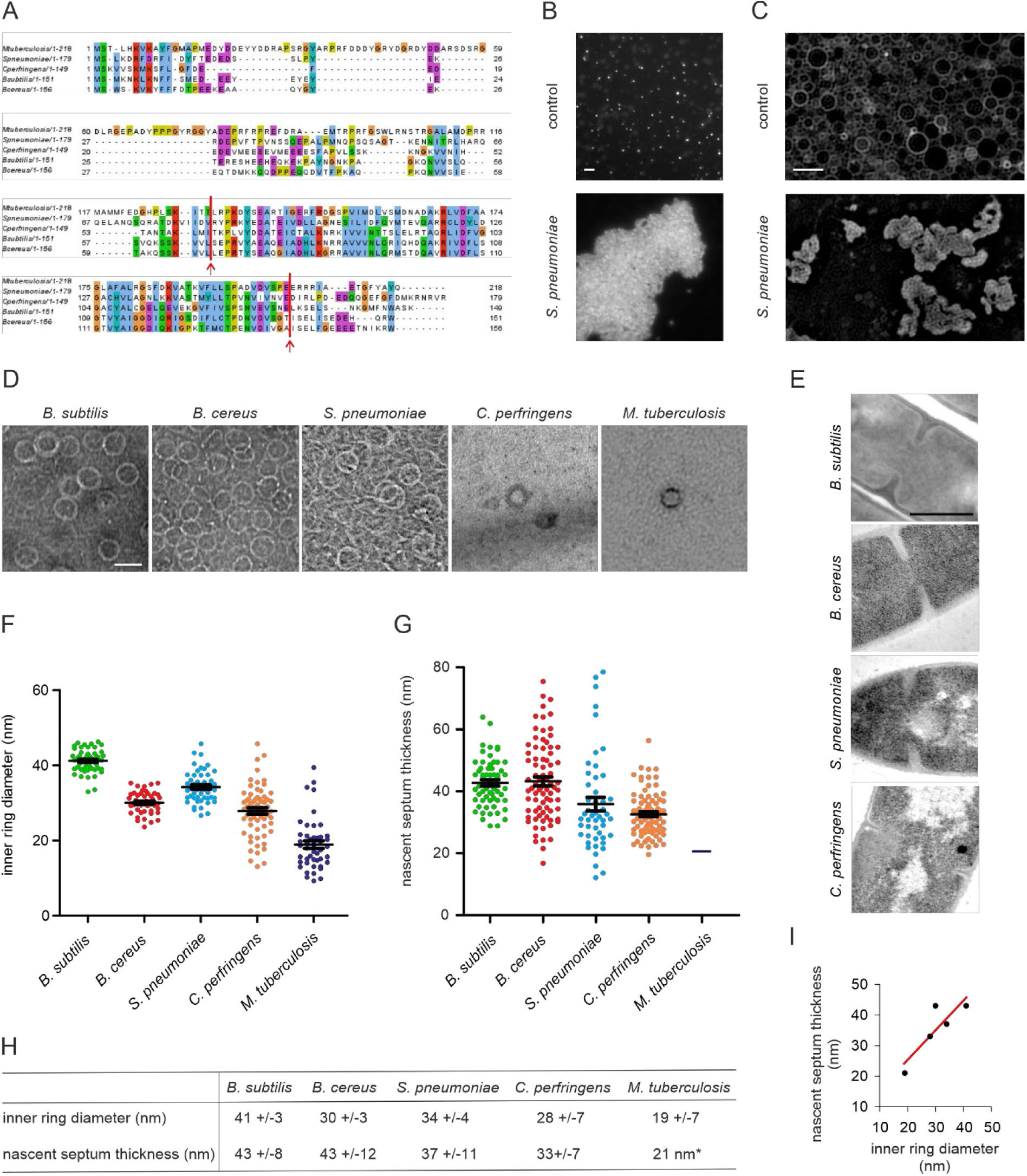
Membrane binding, ring formation and septum thickness. **(A)** Sequence alignment (Clustal Omega) of SepF homologues used in this study. Dark and light grey marks identical and similar amino acids, respectively. Arrows mark the beginning and end of the highly conserved core domain. **(B)** Fluorescence light microscopy image showing the aggregation of small liposomes (∼200 nm) by purified *S. pneumoniae* SepF. Liposomes were stained with Nile red. **(C)** SIM image showing deformation of large liposomes (800 nm) by purified *S. pneumoniae* SepF. Liposomes were stained with mitotracker green. 0.25 µg/ml SepF was mixed with 2 mg/ml liposomes. Scale bars 1 µm. See Fig. S3 for the results with other SepF variants. **(D)** TEM pictures of purified SepF from the different species. Scale bar 50 nm. **(E)** Quantification of the inner SepF ring diameter. Ring diameter was consistently measured as the inner diameter of the SepF ring at its widest point. Black bars show mean with standard error of the mean. A minimum of 45 rings was measured. **(F)** TEM images of nascent cell division septa of the respective species. **(G)** Quantification of the thickness of the nascent septa of the respective organisms. Black bars show mean with standard error of the mean. A minimum of 50 septa was measured. **(H)** Comparison of inner ring diameters with nascent septum thickness. Average +/- standard deviations is indicated. *Data for *M. tuberculosis* were taken from the literature (23). **(I)** Correlation between inner SepF ring diameter and septum thickness (R^2^=0.74).

### Membrane binding

A key property of *B. subtilis* SepF is its capacity to bind to the cell membrane, which is achieved by an N-terminal amphipathic α-helix (amino acids 1-13) (15). This N-terminal amphipathic helix is reasonably conserved in the different SepF variants (Fig. 2A, see Fig. S1 for helical wheel depictions), although the helices differ in their hydrophobicity and hydrophobic moment (Fig. S2). To confirm that the different SepF molecules are able to bind to lipid membranes, the purified proteins were mixed with liposomes. This caused a strong aggregation of liposomes, and deformed them into small vesicles, when observed with high-resolution structured illumination microscopy (SIM). One example, *S. pneumoniae* SepF, is presented in Fig. 2B, C (see Fig. S3 for all variants). Such liposome deformation is a typical characteristic of membrane-interacting amphipathic α-helices (22), and a *B. subtilis* SepF variant without the N-terminal α-helix does not possess this property (15). These results indicate that the membrane binding activity of SepF is conserved.

### Ring formation

Another characteristic of *B. subtilis* SepF is its tendency to polymerize into a curved structure. If this feature is important for the activity of SepF, it should not be restricted to the *B. subtilis* protein. To check this, the purified SepF homologues were spotted on TEM grids and negatively stained with uranyl acetate. Indeed, as shown in Fig. 2D, all SepF variants formed large rings of a remarkably constant size. However, the average ring diameters varied between species, from ∼20 up to ∼40 nm (Fig. 2F).

### Correlation with septum thickness

The different ring diameters provided a first proof of the hypothesis that SepF ring diameter determines septum thickness. To confirm this we performed TEM of the different bacterial species and measured the thickness of nascent cell division septa (Fig. 2E, G). Because of its high pathogenicity we used published data for *M. tuberculosis* (23). As shown in Fig. 2I, there is a clear correlation between inner ring diameters and the thickness of septa in the different species. Only *B. cereus* did not follow this trend, with SepF rings that are clearly smaller than the width of its septa (Fig. 2F-H).

### Core domain defines ring diameter

Proteolytic trimming has shown that the conserved C-terminal domain of *B. subtilis* SepF, spanning amino acids 57 to 151, is sufficient for the formation of rings (15). Therefore, we assumed that this core domain determines ring diameter. To test this, we replaced the conserved core domain of *B. subtilis* SepF with those of the SepF proteins from the other species, maintaining the first 67 and last 13 amino acids of *B. subtilis* SepF. These chimeras were then expressed in *E. coli*, purified, and visualized by TEM using negative staining. All SepF chimeras were able to form rings, and importantly, the ring diameters of these chimeras corresponded very well to the diameters of the original SepF variants (Fig. 3A-D).

**Fig. 3.**
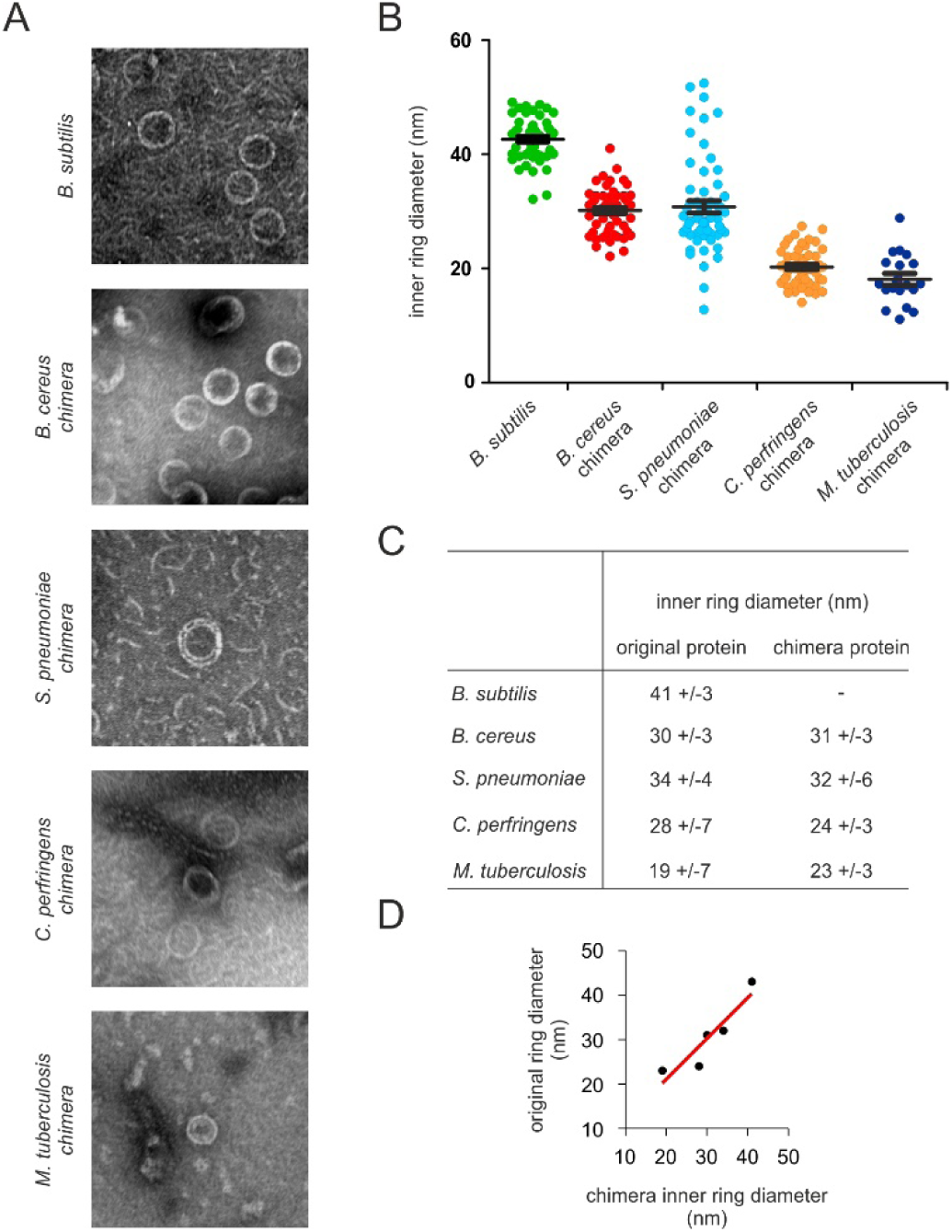
The conserved core domain of SepF determines ring diameter. **(A)** TEM pictures of purified SepF chimera proteins. **(B)** Quantification of ring diameter. Black bars represent mean with standard error of the mean. A minimum of 50 rings was measured. **(C)** Comparison of the inner ring diameters of the original and chimera proteins. Average +/-standard deviation is given. **(D)** Correlation between original and chimera ring diameter (R^2^=0.71).

### Functional chimeras

If it is true that the diameter of SepF rings determines the thickness of septa, then expressing a SepF variant with a smaller diameter in *B. subtilis* should result in thinner septa. It is unlikely that SepF from other species function in *B. subtilis*, since the N- and C-termini of SepF, which are much less conserved, are crucial for its activity (15, 16). Therefore, the chimeras provided a unique opportunity to test this. To determine whether these chimeras are active in *B. subtilis*, they were expressed in a Δ*sepF* strain. *B. subtilis* is one of the few species that can grow without SepF, although this leads to strongly deformed septa (6). Since these deformed septa are lined by the cytoplasmic membrane, this phenotype can be easily observed using high-resolution SIM microscopy by fluorescently labelling the cell membrane (Fig. 4A). Only the *S. pneumoniae* chimera failed to rescue the Δ*sepF* phenotype, while the *B. cereus, C. perfringens,* and *M. tuberculosis* chimeras were all able to restore normal septum formation in the Δ*sepF* background (Fig. 4A). To check that the membrane deformations observed by SIM are indeed indicative of deformed cell wall septa, we confirmed these findings by TEM (Fig. 4B).

**Fig. 4.**
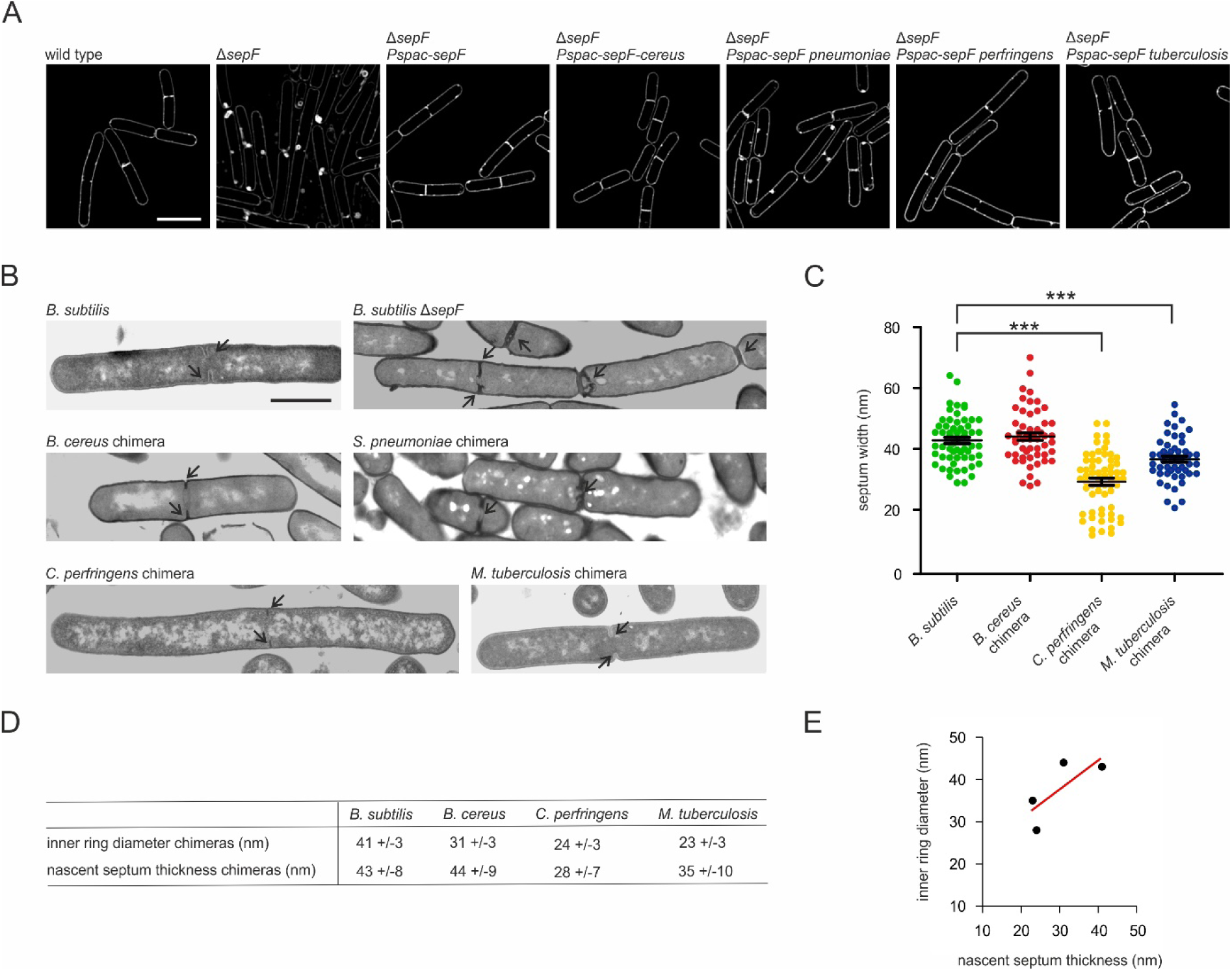
SepF ring diameter determines septum thickness. **(A)** SIM images of *B. subtilis* Δ*sepF* cells expressing different SepF chimeras. The chimeras were expressed from the IPTG-inducible *Pspac* promotor. Cells were grown in the presence of 100 µM IPTG until mid-log phase prior to membrane staining and microscopy. Lack of SepF results in severely deformed septa, which shows as highly fluorescent membrane patches due to membrane invaginations and double membranes. Scale bar 2 µm. **(B)** Representative TEM images of septa of Δ*sepF* strains expressing the different SepF chimeras. Scale bar 1 µm. **(C)** Quantification of the septum thickness of the different strains. Black bars represent mean with standard error of the mean. p values for *C. perfringens* and *M. tuberculosis* chimeras compared to *B. subtilis* are <0.0001. **(D)** Comparison and **(E)** correlation of the inner ring diameter of SepF chimeras and nascent septum thickness of *B. subtilis* Δ*sepF* expressing the chimeras (R^2^=0.57).

The inner ring diameter of the *C. perfringens* SepF chimera (24 nm) is considerably smaller than *B. subtilis* SepF (42 nm). When this chimera was expressed in the Δ*sepF* strain, the septum thickness decreased substantially from 43 to 28 nm (Fig. 4C, D). A similar decrease in septum thickness was observed, when the *M. tuberculosis* SepF chimera (35 nm diameter) was expressed in the Δ*sepF* background (Fig. 4C-E). These results strongly suggests that the SepF ring diameter control septum thickness.

Finally, the diameter of *B. cereus* SepF is approximately 10 nm smaller than *B. subtilis* SepF (Fig. 2F), and so is the diameter of the *B. cereus* SepF chimera (Fig. 3B). However, *B. cereus* has an average septum thickness comparable to that of *B. subtilis* (Fig. 2G). If the core domain of SepF determines septum thickness, it is likely that expression of the *B. cereus* chimera in *B. subtilis* Δ*sepF* results in septa with a thickness comparable to that of wild type *B. subtilis* cells. This was indeed the case (Fig. 4C-E).

## Discussion

In this study we have shown that the membrane binding and ring forming activities are conserved in SepF homologues, that the correlation between ring diameter and septum thickness can be found in different Gram-positive bacteria, and that the septum thickness of *B. subtilis* cells can be reduced by expressing SepF mutants with a smaller diameter. Together these findings strongly suggest that the diameter of curved SepF polymers define septum thickness. The positioning of SepF polymers at the division site cannot be directly observed using high-resolution cryo-electron microscopy due to the electron density of the bacterial cytoplasm and cell wall. However, our findings provide support for a SepF clamp model (Fig. 5). Since the membrane-binding amphipathic α-helix of SepF is located inside the rings, it is likely that SepF polymers do not form rings, but instead form arcs, wrapping the leading edge of nascent septa, on top of which FtsZ polymers bind and align. Since these arcs control the freedom of movement of FtsZ polymers, including the peptidoglycan synthetic apparatus that is linked to them, the diameter of SepF rings will determine the thickness of the septal wall. Many SepF arcs must line the leading edge of the nascent septum to maintain an even thickness. This could be facilitated by the propensity of SepF rings to form stacks (Fig. 1F).

**Fig. 5.**
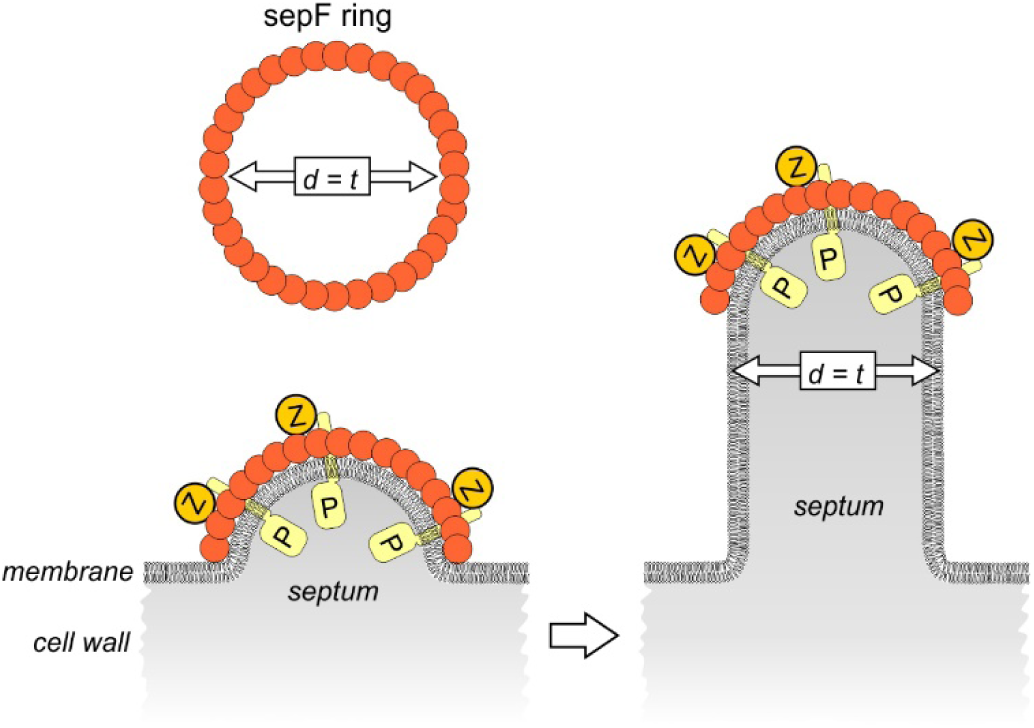
Clamp model illustrating how SepF polymers can control septal thickness. SepF forms large rings with an average inner ring diameter (d) that corresponds to the thickness of septa (t). Crossections of a nascent septum (grey) is depicted. FtsZ polymers (Z) bind to the outside of SepF rings, in a perpendicular fasion. The transmembrane peptidoglycan synthesis machinery (P) is associated with FtsZ polymers. Of note, it is not yet know how these transmembrane proteins are linked to FtsZ polymers in Gram-positive bacteria, and this association might be indirect.

Previous structural work has indicated that SepF rings are built of tightly packed dimers. These dimers are formed by two parallel oriented monomers (15), as a consequence all SepF dimers have the same orientation in a ring. However, FtsZ polymers treadmill in both directions in the Z-ring (26), and it is likely that they bind SepF rings in both polymer orientations. SepF binds to the C-terminal domain of FtsZ (27), which is connected to the main protein domain by a long flexible linker (28). This flexibility might enable FtsZ polymers with opposite orientations to interact with the same SepF ring.

*B. cereus* SepF behaved slightly differently and showed an average inner ring diameter of 30 nm, which is smaller than its average septum thickness of 43 nm. Possibly, *B. cereus* SepF polymers are more flexible in a cellular environment, enabling a wider diameter, or there might be other factors involved in controlling septum diameter in this organism in addition to the physical confinement of SepF polymers. In fact, *B. subtilis* can still make septa, albeit irregular in shape, when SepF is absent, suggesting that other factors provide some control of FtsZ polymers. FtsA, which also forms polymers, might be such factor.

Gram-negative bacteria contain a cell wall that consists of a single-layer of peptidoglycan. The lipoproteins LpoA and LpoB, which reach through the peptidoglycan layer to contact and regulate the cell membrane-anchored peptidoglycan-synthetic enzymes, are thought to regulate peptidoglycan thickness in these species (24). Possibly, this is why Gram-negative bacteria do not need a SepF-like protein for septum thickness control. It should be mentioned that *E. coli* FtsA has been shown to form ring structures *in vitro* as well (25). However, the inner diameter of these rings is ∼15 nm, which is considerably smaller than the width of septa.

Regulation of the shape of septal walls by SepF through its intrinsic curvature is reminiscent of the activity of MreB, which controls the rod shape of many bacteria (29). MreB forms polymers of defined curvature that orientate and move transversely to the long axis of cells, along the direction of greatest membrane curvature (30, 31), thereby guiding the cell wall synthetic machinery in a way that enforces and maintains a rod shape (32, 33). However, unlike SepF, the intrinsic curvature of MreB filaments, which has been estimated to have a diameter of roughly 200 nm (30), does not correlate with the diameter of the cell, which is much larger (30). Thus, the control of septum thickness by a molecular clamp, such as a SepF arc, is a novel concept of how protein polymers can control the shape of growing cell walls.

## Material and Methods

### Strain construction

All strains used in this study are listed in Table S1, plasmids in Table S2, and primers in Table S3. Plasmids for purification of SepF variants were constructed by PCR amplification of *sepF* from the respective organisms, followed by restriction cloning into the pMalC2 plasmid (34), using the *Xba*I and *Sma*I or *Eco*RI restriction sites. pMalC2-based plasmids for purification of chimera proteins and pAPNC-213-kan-based plasmids (35) for integration of *sepF* variants into the *aprE* locus in the *B. subtilis* genome were constructed by Gibson assembly (36). pMalC2-derived purification plasmids were transformed into calcium-competent *E. coli* BL21 (DE3). pAPNC-213-kan-derived integration plasmids were transformed into *B. subtilis* 168 using a standard starvation protocol (37). Deletions of the *sepF* and *ftsA* genes were introduced by transforming the resulting *B. subtilis* strains with chromosomal DNA isolated from YK204 (*sepF::spc*) (38), BFA2863 (*sepF::ery*) (6), and TB94 (*ftsA::cat*), respectively. TB94 was created by transformation of *B. subtilis* 168 with chromosomal DNA of strain MD133 *ftsA::cat aprE::kan Pspac-ftsA* (38) followed by selection on chloramphenicol resistance but kanamycin sensitivity.

### Protein purification

*E. coli* BL21 (DE3) strains carrying pMalC2 plasmids for purification of SepF variants were grown overnight in LB containing 100 µg/ml ampicillin at 37 °C under continuous shaking. Cultures were diluted 1:100 and allowed to grow until an OD_600_ of 0.4 in the presence of ampicillin. Expression of maltose-binding protein (MBP)-tagged SepF proteins was induced by addition of 0.5 mM IPTG. Cells were harvested by centrifugation after 4 h of induction and subsequently washed in phosphate-buffered saline supplemented with 1 mM phenylmethylsulfonyl fluoride to prevent protein degradation. Cell pellets were flash-frozen in liquid nitrogen and stored at −80 °C until further use. Cell pellets were resuspended in buffer AF (50 mM Tris-HCL, pH 7.4, 200 mM KCl, 5 mM EDTA, 0.5 mM DTT) supplemented with one Complete mini protease inhibitor tablet (Roche) and disrupted by French Press. Cell debris was removed by ultracentrifugation at 31,000 x g, and the resulting supernatant was filtered through 0.2 µm filter membranes prior to loading onto a Tricorn 10/20 column (GE Healthcare) packed with 2 ml amylose resin (New England Biolabs) equilibrated with buffer AF. After loading, the column was washed with 5 column volumes buffer AF, followed by buffer BF (50 mM Tris-HCl, pH 7.4) until the baseline was stable. MBP-tagged SepF variants were eluted with buffer BF containing 10 mM maltose. Appropriate fractions were pooled and digested with factor Xa protease (New England Biolabs) in the presence of 2 mM CaCl_2_ at 4 °C overnight. *B. cereus* SepF was soluble while the others precipitated after cleavage (*B. subtilis, S. pneumoniae, C. perfringens, M. tuberculosis,* all chimera proteins) due to presence of calcium. Insoluble proteins were separated from soluble MBP and factor Xa by centrifugation at 10,000 x g. Pellets containing pure SepF were dissolved in buffer BF without CaCl_2_, flash-frozen in liquid nitrogen and stored at −80 °C until further use. SepF variants that were soluble after factor Xa cleavage were separated from MBP and Factor Xa protease by ion exchange chromatography. To this end, digested samples were loaded onto a 1 ml HiTrap Q column (GE Healthcare) equilibrated with buffer BF. The column was washed with buffer BF until the baseline was stable, followed by washing with 7.5% buffer CF (50 mM Tris-HCl, pH 7.4, 1 M KCl) resulting in elution of MBP, and washing with 17.5% buffer CF resulting in elution of factor Xa. Pure SepF was eluted with 50% buffer CF, flash-frozen in liquid nitrogen, and stored at −80 °C.

### Atomic force microscopy

Purified *B. subtilis* SepF was diluted 1:10 in buffer BF and added onto a freshly cleaved highly oriented pyrolytic graphite (HOPG) surface (Agar Scientific) and allowed to settle for 20 min at room temperature prior to imaging in buffer solution (39). Images were taken with an atomic force microscope from Nanotec Electronica (Tres Cantos) operated in jumping mode at room temperature. Olympus OMCL-RC800PSA rectangular, silicon-nitride cantilevers with a nominal tip radius of 15 nm and a nominal spring constant of 0.05 N/m were used. Images were processed and analyzed with WSxM software (40).

### Fluorescence light microscopy of liposomes

Liposomes were prepared from *E. coli* polar lipid extract (Avanti Polar Lipids) as described previously (3). Liposomes were extruded through 0.2 µm filters. Samples were stained with 1 µg/ml Nile red, spotted on 1.2% agarose films, covered with poly-dopamine-coated cover slips (41), and immediately imaged with a Nikon Eclipse Ti equipped with a CFI Plan Apochromat DM 100x oil objective, an Intensilight HG 130 W lamp, a C11440-22CU Hamamatsu ORCA camera, and NIS elements software. Images were analyzed using ImageJ (National Institutes of Health).

### Structured illumination (SIM) microscopy

Liposomes for SIM were prepared from *E. coli* polar lipid extract (Avanti Polar Lipids) as described previously (3). Liposomes were extruded through 0.8 µm filters to obtain large enough vesicles. 0.25 mg/ml of the respective SepF variants was mixed with 2 mg/ml liposomes in SepF binding buffer. Samples were stained with 0.5 µg/ml mitotracker green, spotted on 1.2% agarose films, covered with poly-dopamine-coated cover slips (41), and immediately imaged with a Nikon Eclipse Ti N-SIM E microscope setup equipped with a CFI SR Apochromat TIRF 100x oil objective (NA1.49), a LU-N3-SIM laser unit, an Orca-Flash 4.0 sCMOS camera (Hamamatsu Photonics K.K.), and NIS elements Ar software. SIM microscopy of bacteria was performed by staining cells with 0.5 µg/ml mitotracker green for 1 min, spotted on a thin film of 1.2% agarose (21).

### Transmission electron microscopy of proteins

Protein samples were spotted on glow-discharged 200 mesh formvar/carbon-coated copper grids (Agar Scientific) and incubated for 1 min at room temperature. Excess liquid was removed with paper tissue and samples were negatively stained by adding 100 µl 2% uranyl acetate drop by drop. Excess staining solution was removed with paper tissue and samples were allowed to air dry. Samples were examined with a JEOL1010 at 60 kV.

### Growth conditions for microscopy

*B. cereus, S. pneumoniae*, and *C. perfringens* were grown on tryptic soy agar plates containing 5 % sheep blood (BioMerieux). After three days of aerobic (*B. cereus* and *S. pneumoniae*) or anaerobic incubation (*C. perfringens*) at 37 °C, colonies were transferred to fresh plates and incubated for another 48 h prior to suspension in phosphate-buffered saline and preparation for electron microscopy. *B. subtilis* strains were grown in Luria-Bertani (LB) broth at 37 °C under steady agitation. The medium was supplemented with 0.1 mM IPTG to induce expression of SepF variants, where appropriate. It is important not to use more IPTG since SepF overproduction causes membrane deformations that obscure septa (42). Overnight cultures were grown with appropriate antibiotic concentrations (100 µg/ml spectinomycin, 10 µg/ml chloramphenicol, 5 µg/ml kanamycin, 1 µg/ml erythromycin), where necessary. Cells were grown until exponential phase (OD_600_ = 0.4) prior to microscopy.

### Transmission electron microscopy of bacteria

Electron microscopy of bacteria was performed according to van Wezel *at al.* (43) (*B. cereus*, *S. pneumoniae*, *C. perfringens)* or to a novel method that uses immobilization of bacterial cells in one plane on an agarose layer prior to fixation and embedding (44) (*B. subtilis*). Samples were examined with a JEOL 1010 at an electron voltage of 60 kV.

## Acknowledgements

We gratefully acknowledge Zehui Zhang and Daniel Antwi-berko for help with sample preparation, Wiep Klaas Smits for help with growing strains, Marien P. Dekker and Jan R. T. van Weering for support with TEM, and Gaurav Dugar for critically reading the manuscript. Electron microscopy was performed at the VU/VUMC EM facility, supported by the Netherlands Organization for Scientific Research (NWO, middelgroot 91111009). This work was financially supported by the Netherlands Organization for Scientific Research (NWO, STW-Vici 12128 to LWH and NWO-Vidi to WHR). MW was supported by a postdoc stipend from the Amsterdam Infection and Immunity Institute.

## Author contributions

MW planned and performed experiments and wrote the paper. INCG constructed strains and performed experiments. YG constructed strains. JW and MGM performed experiments. GJLW and WHR provided tools and planned experiments. LWH designed the study and wrote the paper.

## Supplementary Information

**Table S1.** Strains used in this study.

| **strains** | **genotype** | **reference** |
| --- | --- | --- |
| *B. subtilis* 168 | *trpC2* | (1) |
| *B. subtilis* TB92 | *trpC2 sepF::spc* (YK204 in 168CA) | (2) |
| *B. subtilis* TB94 | *trpC2 ftsA::cat* (MD133 in 168CA) | (2) |
| *B. subtilis* LH28 | *trpC2 ezrA::cat* | (3) |
| *B. subtilis* BFA2863 | *trpC2 sepF::ery* | (4) |
| *B. subtilis* GYQ457 | *trpC2 aprE::kan Pspac-sepF(subtilis-cereus)* | this study |
| *B. subtilis* GYQ475 | *trpC2 aprE::kan Pspac-sepF(subtilis-pneumoniae)* | this study |
| *B. subtilis* GYQ544 | *trpC2 aprE::kan Pspac-sepF(subtilis-perfringens)* | this study |
| *B. subtilis* GYQ474 | *trpC2 aprE::kan Pspac-sepF(subtilis-tuberculosis)* | this study |
| *B. subtilis* MW18 | *trpC2 aprE::kan Pspac-sepF(WT) sepF::spc* | this study |
| *B. subtilis* GYQ458 | *trpC2 aprE::kan Pspac-sepF(subtilis-cereus)* | this study |
| *B. subtilis* MW89 | *trpC2 aprE::kan Pspac-sepF(subtilis-pneumoniae)* | this study |
| *B. subtilis* MW92 | *trpC2 aprE::kan Pspac-sepF(subtilis-perfringens)* | this study |
| *B. subtilis* MW86 | *trpC2 aprE::kan Pspac-sepF(subtilis-tuberculosis)* | this study |
| *E. coli* BL21(DE03) pNC1 | *pMalC2-sepF(coelicolor3)* | this study |
| *E. coli* BL21(DE03) pNC2 | *pMalC2-sepF(cereus)* | this study |
| *E. coli* BL21(DE03) pNC4 | *pMalC2-sepF(tuberculosis)* | this study |
| *E. coli* BL21(DE03) pNC5 | *pMalC2sepF(coelicolor2)* | this study |
| *E. coli* BL21(DE03) pNC6 | *pMalC2-sepF(coelicolor1)* | this study |
| *E. coli* BL21(DE03) pNC7 | *pMalC2-sepF(perfringens)* | this study |
| *E. coli* BL21(DE03) pNC9 | *pMalC2-sepF(pneumoniae)* | this study |
| *E. coli* BL21(DE03) pNC12 | *pMalC2-sepF(subtilis)* | (5) |
| *E. coli* BL21(DE03) pYQ62 | *pMalC2-sepF(subtilis-cereus)* | this study |
| *E. coli* BL21(DE03) pYQ94 | *pMalC2-sepF(subtilis-tuberculosis)* | this study |
| *E. coli* BL21(DE03) pYQ95 | *pMalC2-sepF(subtilis-perfringens)* | this study |
| *E. coli* BL21(DE03) pYQ96 | *pMalC2-sepF(subtilis-pneumoniae)* | this study |
| *B. cereus* ATCC14579 |  | (6) |
| *S. pneumoniae* ATCC6305 |  | (7, 8) |
| *C. perfringens* ATCC13124 |  | (9) |

**Table S2.** Plasmids used in this study.

| **plasmid** | **genotype** | **reference** |
| --- | --- | --- |
| pMW8 | *pAPNC-213-kan-Pspac-sepF(WT)* | this study |
| pYQ90 | *pAPNC-213-kan-Pspac-sepF(subtilis-cereus)* | this study |
| pYQ97 | *pAPNC-213-kan-Pspac-sepF(subtilis-tuberculosis)* | this study |
| pYQ98 | *pAPNC-213-kan-Pspac-sepF(subtilis-perfringens)* | this study |
| pYQ99 | *pAPNC-213-kan-Pspac-sepF(subtilis-pneumoniae)* | this study |
| pNC1 | *pMalC2-sepF(coelicolor3)* | this study |
| pNC2 | *pMalC2-sepF(cereus)* | this study |
| pNC4 | *pMalC2-sepF(tuberculosis)* | this study |
| pNC5 | *pMalC2sepF(coelicolor2)* | this study |
| pNC6 | *pMalC2-sepF(coelicolor1)* | this study |
| pNC7 | *pMalC2-sepF(perfringens)* | this study |
| pNC9 | *pMalC2-sepF(pneumoniae)* | this study |
| pNC12 | *pMalC2-sepF(subtilis)* | (5) |
| pYQ62 | *pMalC2-sepF(subtilis-cereus)* | this study |
| pYQ94 | *pMalC2-sepF(subtilis-tuberculosis)* | this study |
| pYQ95 | *pMalC2-sepF(subtilis-perfringens)* | this study |
| pYQ96 | *pMalC2-sepF(subtilis-pneumoniae)* | this study |

**Table S3.**
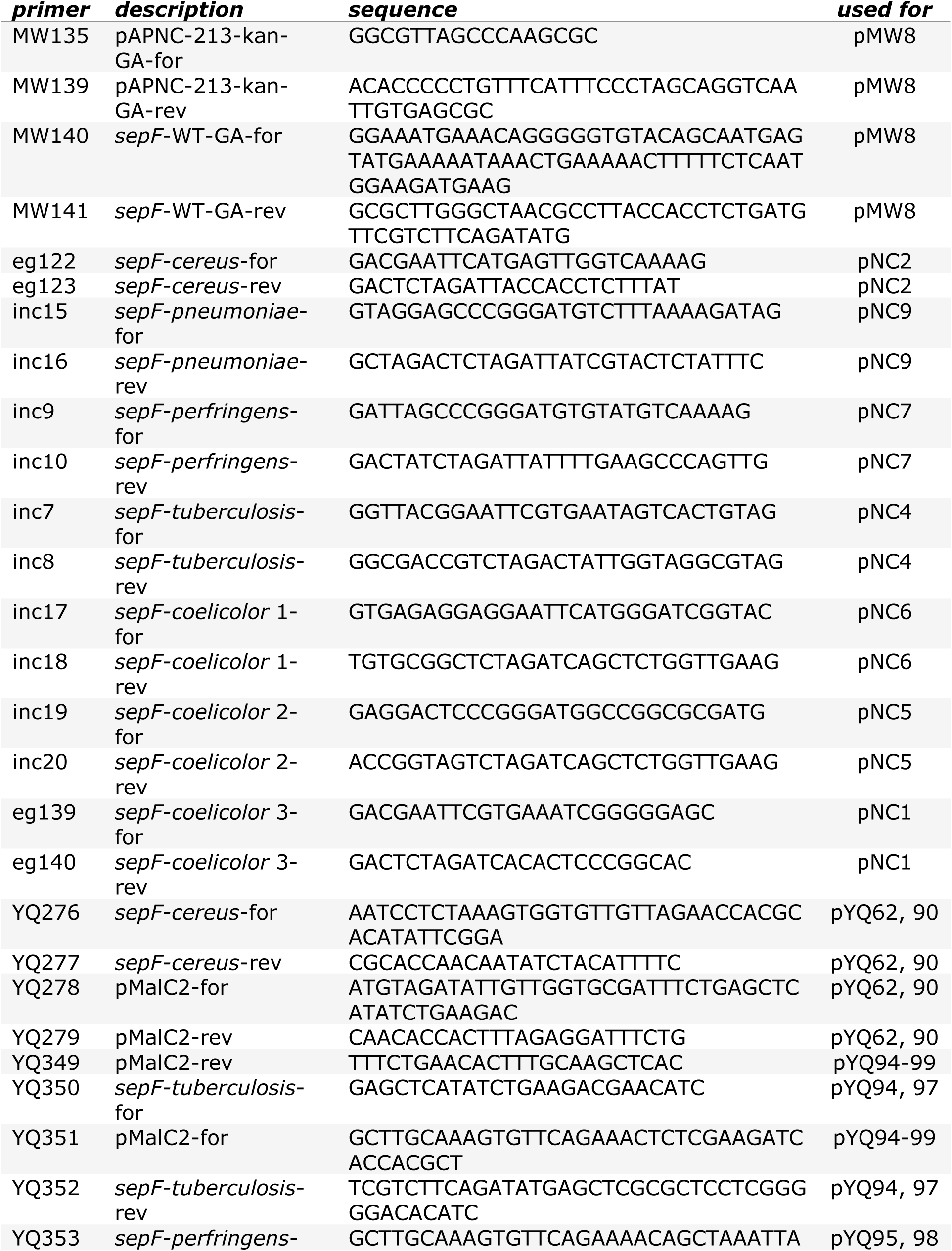
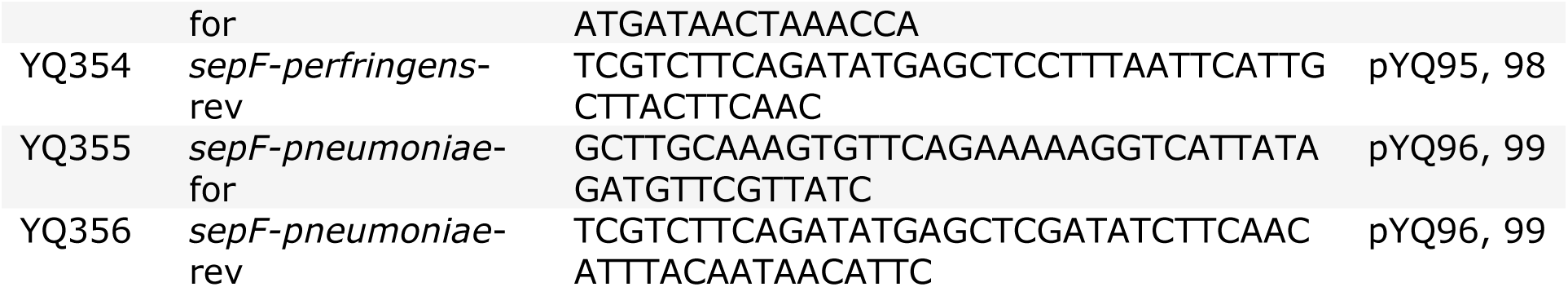
Primers used in this study.

**Fig. S1.**
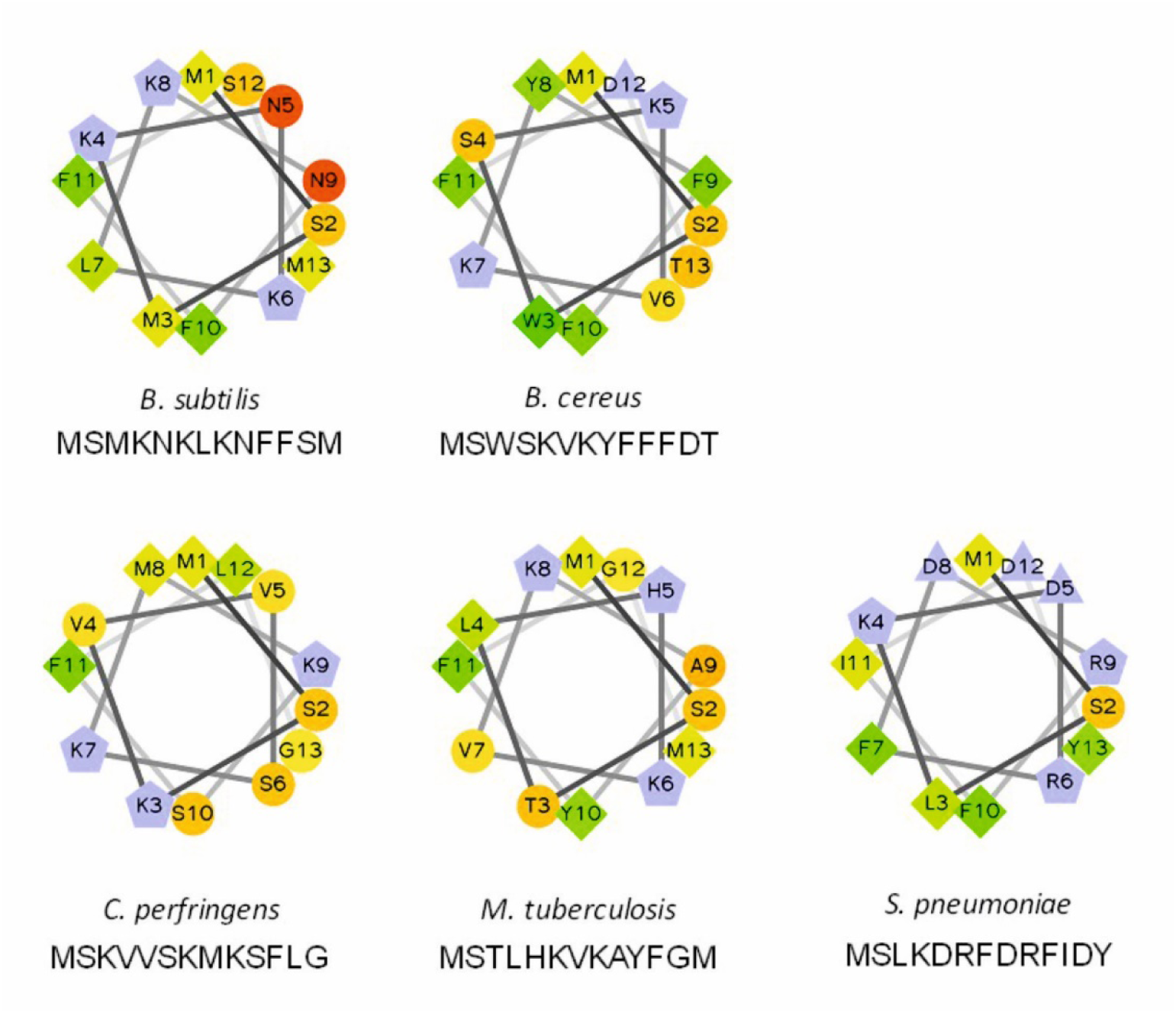
Comparison of the amphipathic helices of SepF variants. Helical wheel projections (RZ lab) show the first 13 amino acids of the different proteins. Hydrophilic residues are represented as circles, hydrophobic residues as diamonds, negatively charged as triangles, and positively charged as pentagons. Hydrophobicity is color coded with green indicating the highest and yellow the lowest (zero) hydrophobicity. Hydrophilic residues are depicted in red with the amount of red decreasing to orange proportionally to the hydrophilicity. Potentially charged residues are depicted in light blue.

**Fig. S2.**
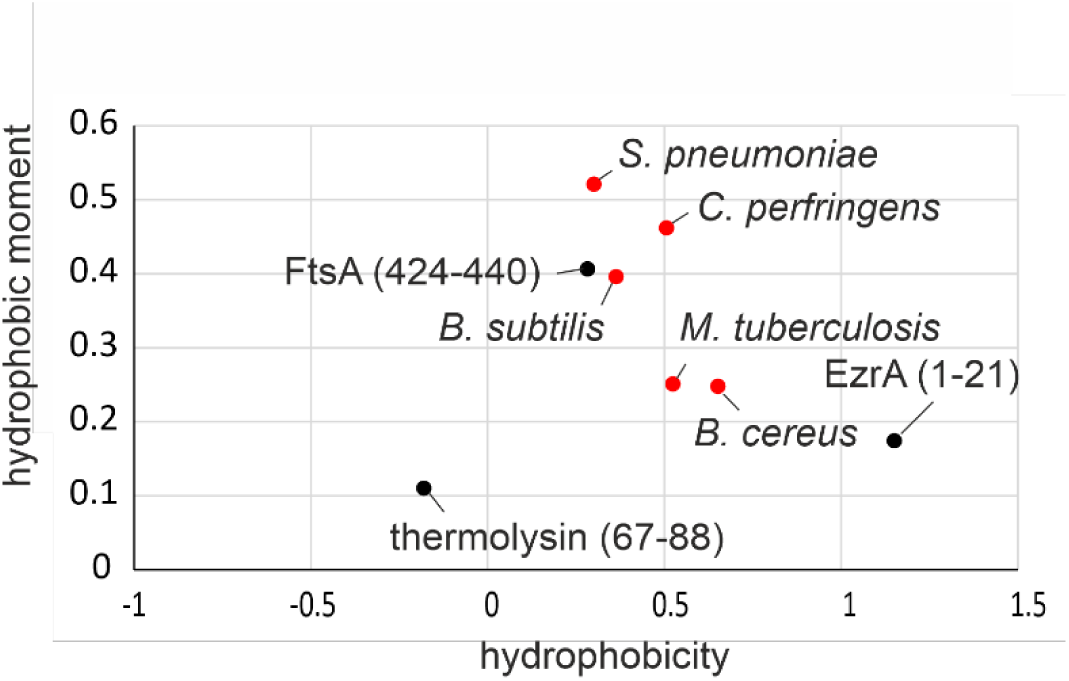
Hydrophobic moment plot of the putative N-terminal α-helices of SepF variants. High H values indicate strong hydrophobicity. The hydrophobic moment (µH) is a measure for the amphipathicity of an α-helix (high values indicating strong amphipathicity). The plot of H and µH indicates the probability of an α-helix to reside in the globular or surface-exposed domains of a protein and can be used to predict transmembrane or amphipathic membrane-binding helices. Black dots indicate known globular (*Bacillus thermoproteolyticus* thermolysin, aa 67-88), transmembrane (*B. subtilis* EzrA, aa 1-21) and amphipathic (*B. subtilis* FtsA, aa 424-440) helices for comparison. N-terminal α-helices of SepF homologues are depicted in red.

**Fig. S3.**
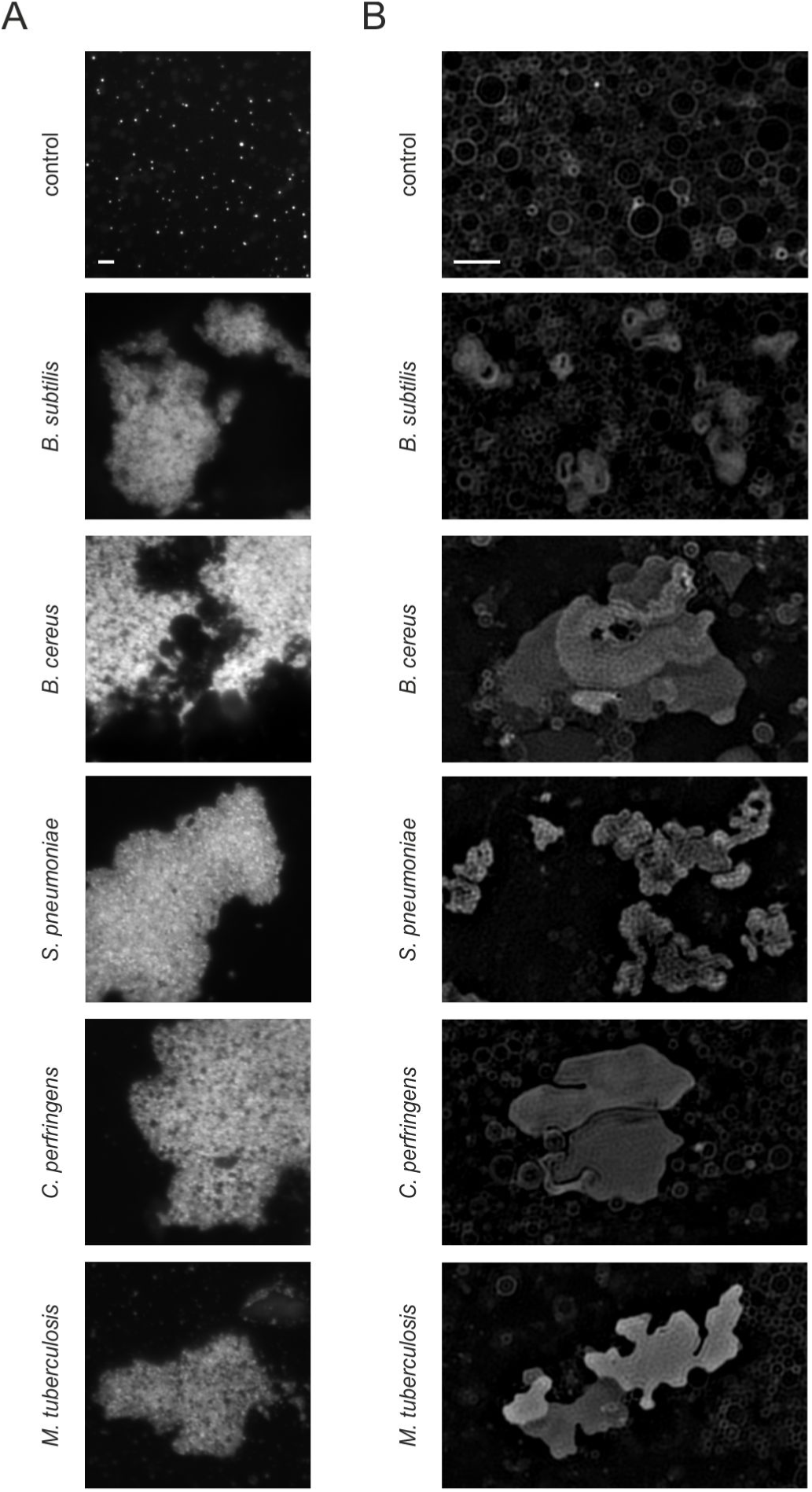
Liposome aggregation and deformation by SepF variants. **(A)** Fluorescence light microscopy images of small liposomes (200 nm) stained with Nile red. **(B)** SIM images of large liposomes (800 nm) stained with mitotracker green. 0.25 µg/ml of the different SepF proteins (species names indicated) were mixed with 2 mg/ml liposomes prepared from *E. coli* polar lipid extract. Scale bars 1 µm.

## References

1. J. Lutkenhaus, S. Pichoff, S. Du, Bacterial cytokinesis: From Z ring to divisome. Cytoskeleton 69, 778–790 (2012).

2. P. Gamba, J. W. Veening, N. J. Saunders, L. W. Hamoen, R. A. Daniel, Two-step assembly dynamics of the Bacillus subtilis divisome. J. Bacteriol. 191, 4186–4194 (2009).

3. H. Strahl, L. W. Hamoen, Membrane potential is important for bacterial cell division. Proc. Natl. Acad. Sci. U. S. A. 107, 12281–12286 (2010).

4. S. Pichoff, J. Lutkenhaus, Tethering the Z ring to the membrane through a conserved membrane targeting sequence in FtsA. Mol. Microbiol. 55, 1722–1734 (2005).

5. J. Errington, R. A. Daniel, D.-J. Scheffers, Cytokinesis in bacteria. Microbiol. Mol. Biol. Rev. 67, 52–65, table of contents (2003).

6. L. W. Hamoen, J.-C. Meile, W. De Jong, P. Noirot, J. Errington, SepF, a novel FtsZ-interacting protein required for a late step in cell division. Mol. Microbiol. 59, 989–999 (2006).

7. F. J. Gueiros-Filho, R. Losick, A widely conserved bacterial cell division protein that promotes assembly of the tubulin-like protein FtsZ. Genes Dev. 16, 2544–2556 (2002).

8. P. A. Levin, I. G. Kurtser, A. D. Grossman, Identification and characterization of a negative regulator of FtsZ ring formation in Bacillus subtilis. Proc. Natl. Acad. Sci. U. S. A. 96, 9642–9647 (1999).

9. C. A. Hale, P. A. de Boer, Direct binding of FtsZ to ZipA, an essential component of the septal ring structure that mediates cell division in E. coli. Cell 88, 175–185 (1997).

10. A. Taguchi, et al., FtsW is a peptidoglycan polymerase that is functional only in complex with its cognate penicillin-binding protein. Nat. Microbiol. 4, 587–594 (2019).

11. R. A. Daniel, E. J. Harry, J. Errington, Role of penicillin-binding protein PBP 2B in assembly and functioning of the division machinery of Bacillus subtilis. Mol. Microbiol. 35, 299–311 (2000).

12. L. Wang, M. K. Khattar, W. D. Donachie, J. Lutkenhaus, FtsI and FtsW are localized to the septum in Escherichia coli. J. Bacteriol. 180, 2810–2816 (1998).

13. R. A. Daniel, M.-F. Noirot-Gros, P. Noirot, J. Errington, Multiple interactions between the transmembrane division proteins of Bacillus subtilis and the role of FtsL instability in divisome assembly. J. Bacteriol. 188, 7396–7404 (2006).

14. N. Buddelmeijer, J. Beckwith, A complex of the Escherichia coli cell division proteins FtsL, FtsB and FtsQ forms independently of its localization to the septal region. Mol. Microbiol. 52, 1315–1327 (2004).

15. R. Duman, et al., Structural and genetic analyses reveal the protein SepF as a new membrane anchor for the Z ring. Proc. Natl. Acad. Sci. U. S. A. 110, E4601–10 (2013).

16. M. E. Gündoğdu, et al., Large ring polymers align FtsZ polymers for normal septum formation. EMBO J. 30, 617–626 (2011).

17. A. Kotiranta, K. Lounatmaa, M. Haapasalo, Epidemiology and pathogenesis of Bacillus cereus infections. Microbes Infect. 2, 189–198 (2000).

18. E. J. Bottone, Bacillus cereus, a volatile human pathogen. Clin. Microbiol. Rev. 23, 382–398 (2010).

19. K. S. Antonation, et al., Bacillus cereus Biovar Anthracis Causing Anthrax in Sub-Saharan Africa—Chromosomal Monophyly and Broad Geographic Distribution. PLoS Negl. Trop. Dis. 10, e0004923 (2016).

20. A. R. Hoffmaster, et al., Identification of anthrax toxin genes in a Bacillus cereus associated with an illness resembling inhalation anthrax. Proc. Natl. Acad. Sci. 101, 8449–8454 (2004).

21. C. Hoffmann, et al., Persistent anthrax as a major driver of wildlife mortality in a tropical rainforest. Nature 548, 82 (2017).

22. W. A. Prinz, J. E. Hinshaw, Membrane-bending proteins. Crit. Rev. Biochem. Mol. Biol. 44, 278–291 (2009).

23. A. A. Velayati, P. Farnia, Altas of Mycobacterium tuberculosis, 1st Editio (Academic Press, 2016).

24. A. Typas, et al., Regulation of peptidoglycan synthesis by outer-membrane proteins. Cell 143, 1097–1109 (2010).

25. M. Krupka, et al., Escherichia coli FtsA forms lipid-bound minirings that antagonize lateral interactions between FtsZ protofilaments. Nat. Commun. 8, 15957 (2017).

26. A. W. Bisson-Filho, et al., Treadmilling by FtsZ filaments drives peptidoglycan synthesis and bacterial cell division. Science 355, 739–743 (2017).

27. E. Krol, et al., Bacillus subtilis SepF binds to the C-terminus of FtsZ. PLoS One 7, e43293 (2012).

28. P. J. Buske, A. Mittal, R. V Pappu, P. A. Levin, An intrinsically disordered linker plays a critical role in bacterial cell division. Semin. Cell Dev. Biol. 37, 3–10 (2015).

29. L. J. Jones, R. Carballido-Lopez, J. Errington, Control of cell shape in bacteria: helical, actin-like filaments in Bacillus subtilis. Cell 104, 913–922 (2001).

30. S. Hussain, et al., MreB filaments align along greatest principal membrane curvature to orient cell wall synthesis. Elife 7, e32471 (2018).

31. T. S. Ursell, et al., Rod-like bacterial shape is maintained by feedback between cell curvature and cytoskeletal localization. Proc. Natl. Acad. Sci. U. S. A. 111, E1025–34 (2014).

32. J. Dominguez-Escobar, et al., Processive movement of MreB-associated cell wall biosynthetic complexes in bacteria. Science 333, 225–228 (2011).

33. E. C. Garner, et al., Coupled, circumferential motions of the cell wall synthesis machinery and MreB filaments in B. subtilis. Science 333, 222–225 (2011).

34. P. Riggs, “Expression and Purification of Maltose-Binding Protein Fusions” in Current Protocols in Molecular Biology, (John Wiley & Sons, Inc., 2001) https:/doi.org/10.1002/0471142727.mb1606s28.

35. M. Yoshimura, T. Oshima, N. Ogasawara, Involvement of the YneS/YgiH and PlsX proteins in phospholipid biosynthesis in both Bacillus subtilis and Escherichia coli. BMC Microbiol. 7, 69 (2007).

36. D. G. Gibson, et al., Enzymatic assembly of DNA molecules up to several hundred kilobases. Nat Meth 6, 343–345 (2009).

37. P. M. Hauser, D. Karamata, A rapid and simple method for Bacillus subtilis transformation on solid media. Microbiology 140, 1613–1617 (1994).

38. S. Ishikawa, Y. Kawai, K. Hiramatsu, M. Kuwano, N. Ogasawara, A new FtsZ-interacting protein, YlmF, complements the activity of FtsA during progression of cell division in Bacillus subtilis. Mol. Microbiol. 60, 1364–1380 (2006).

39. K. Heinze, et al., Protein Nanocontainers from Nonviral Origin: Testing the Mechanics of Artificial and Natural Protein Cages by AFM. J. Phys. Chem. B 120, 5945–5952 (2016).

40. I. Horcas, et al., WSXM: A software for scanning probe microscopy and a tool for nanotechnology. Rev. Sci. Instrum. 78, 13705 (2007).

41. J. D. te Winkel, D. A. Gray, K. H. Seistrup, L. W. Hamoen, H. Strahl, Analysis of antimicrobial-triggered membrane depolarisation using voltage sensitive dyes. Front. Cell Dev. Biol. 4, 29 (2016).

42. Y. Gao, M. Wenzel, M. J. Jonker, L. W. Hamoen, Free SepF interferes with recruitment of late cell division proteins. Sci. Rep. 7, 16928 (2017).

43. G. P. van Wezel, et al., ssgA is essential for sporulation of Streptomyces coelicolor A3(2) and affects hyphal development by stimulating septum formation. J. Bacteriol. 182, 5653–5662 (2000).

44. D. Saeloh, et al., The novel antibiotic rhodomyrtone traps membrane proteins in vesicles with increased fluidity. PLoS Pathog. 14, e1006876 (2018).

## References

1. Anagnostopoulos C, Spizizen J (1960) Requirements for transformation in Bacillus subtilis. J Bacteriol 81(5):741–746.

2. Ishikawa S, Kawai Y, Hiramatsu K, Kuwano M, Ogasawara N (2006) A new FtsZ-interacting protein, YlmF, complements the activity of FtsA during progression of cell division in Bacillus subtilis. Mol Microbiol 60(6):1364–1380.

3. Gamba P, Rietkotter E, Daniel RA, Hamoen LW (2015) Tetracycline hypersensitivity of an ezrA mutant links GalE and TseB (YpmB) to cell division. Front Microbiol 6:346.

4. Hamoen LW, Meile J-C, De Jong W, Noirot P, Errington J (2006) SepF, a novel FtsZ-interacting protein required for a late step in cell division. Mol Microbiol 59(3):989–999.

5. Duman R, et al. (2013) Structural and genetic analyses reveal the protein SepF as a new membrane anchor for the Z ring. Proc Natl Acad Sci U S A 110(48):E4601–10.

6. Frankland GC, Frankland PF (1887) XI. Studies on some new micro-organisms obtained from air. Philos Trans R Soc London 178:257–287.

7. Public Health Weekly Reports for APRIL 14, 1944 (1944) Public Heal reports (Washington, DC 1896) 59(15):485–512.

8. Public Health Weekly Reports for APRIL 7, 1944 (1944) Public Heal reports (Washington, DC 1896) 59(14):449–484.

9. Eastoe JE, Long JE (1959) The effect of nisin on the growth of cels and spores of Clostridium welchii in gelatine. J Appl Bacteriol 22(1):1–7.

